# Double insertion of transposable elements provides a substrate for the evolution of satellite DNA

**DOI:** 10.1101/158386

**Authors:** Michael P. McGurk, Daniel A. Barbash

## Abstract

Eukaryotic genomes are replete with repeated sequences, in the form of transposable elements (TEs) dispersed across the genome or as satellite arrays, large stretches of tandemly repeated sequence. Many satellites clearly originated as TEs, but it is unclear how mobile genetic parasites can transform into megabase-sized tandem arrays. Comprehensive population genomic sampling is needed to determine the frequency and generative mechanisms of tandem TEs, at all stages from their initial formation to their subsequent expansion and maintenance as satellites. The best available population resources, short-read DNA sequences, are often considered to be of limited utility for analyzing repetitive DNA due to the challenge of mapping individual repeats to unique genomic locations. Here we develop a new pipeline called ConTExt which demonstrates that paired-end Illumina data can be successfully leveraged to identify a wide range of structural variation within repetitive sequence, including tandem elements. Analyzing 85 genomes from five populations of *Drosophila melanogaster* we discover that TEs commonly form tandem dimers. Our results further suggest that insertion site preference is the major mechanism by which dimers arise and that, consequently, dimers form rapidly during periods of active transposition. This abundance of TE dimers has the potential to provide source material for future expansion into satellite arrays, and we discover one such copy number expansion of the DNA transposon *Hobo* to ~16 tandem copies in a single line. The very process that defines TEs —transposition— thus regularly generates sequences from which new satellites can arise.

## Introduction

Eukaryotic genomes are inundated with two types of repetitive sequences: transposable elements (TEs), which are dispersed by a variety of transposition mechanisms, and satellite sequences, which are tandemly repeated sequences that expand, contract, and are homogenized by recombination events. Both types of repeats are enriched near the telomeres and in the heterochromatin surrounding the centromeres, likely because the low frequency of recombination typical of heterochromatin permits their persistence (Charlesworth et al., 1986).

The essential roles played by telomeres and centromeres in genome integrity and chromosome segregation suggest that some repetitive sequences are of functional significance. Examples supporting functional roles for repetitive sequences mostly follow from observations of phenotypes associated with repeat variation. Contractions of the human subtelomeric satellite D4Z4 cause facioscapulohumeral muscular dystrophy by altering the chromatin state of nearby genes (Zeng et al., 2009). Sequence variation in a human centromeric satellite is associated with mitotic segregation errors resulting in aneuploidy (Aldrup-MacDonald et al., 2016). Variants of the mostly repetitive *Drosophila melanogaster* Y chromosome have global impacts on gene expression, possibly by titrating chromatin binding factors (Francisco and Lemos, 2014). Satellites can also engage in meiotic drive and gamete competition (Fishman and Saunders, 2008) (Hardy et al., 1984) (Larracuente, 2014), selfish processes whereby alleles bias meiotic segregation or gamete survival to transmit at frequencies greater than Mendelian expectations. Finally, the structural importance of constitutive heterochromatin means that changes in repeat composition between species can cause reproductive barriers (Ferree and Barbash, 2009).

Despite the potential consequences of satellite variation, many satellite sequences turnover rapidly between closely related species (Lohe and Roberts, 2000). Partially explaning this is the potential of satellite sequence to recombine out of register, termed unequal exchange. Neutral evolution by unequal exchange leads to 1) dramatic changes in copy number from relatively few exchange events and 2) the eventual contraction of the array to a single repeat unit (Charlesworth et al., 1986). The long-term persistence of some conserved satellites (Strachan et al., 1982) may therefore reflect functional importance. Given their ubiquity, however, unless all satellites are functional, mechanisms to generate new satellites must exist to counter the inevitable loss of neutrally evolving ones (Charlesworth et al., 1986).

Models of satellite evolution suggest two stages in the emergence of new satellites: 1) amplification processes generate small tandem sequences, and 2) some of these sequences expand to large arrays by unequal exchange (Stephan and Cho, 1994). Thus, any process that generates sequence upon which unequal exchange can act is a potential source of new satellites. Simple satellites, for example, can readily arise by polymerase slippage and subsequent copy number expansion. These simple satellites can transition to more complex satellite types by the interplay of unequal exchange and mutations (Prosser et al., 1986; Stephan and Cho, 1994).

More enigmatic mechanisms to generate new satellites also exist. TEs are well-known culprits in causing mutations and gene duplication events, but they are also found as tandem arrays in many species, including as centromeric satellites (Meštrović et al., 2015). The easiest to understand are satellites derived from TEs with intrinsic repeats, such as long terminal repeats (LTRs) and tandemly repeated regulatory elements, which provide substrates for expansion by unequal exchange (Fig 1A) (Dias et al., 2014; Gong et al., 2012; Ke and Voytas, 1997; Macas et al., 2009; Zhang et al., 2014).

**Figure 1:**
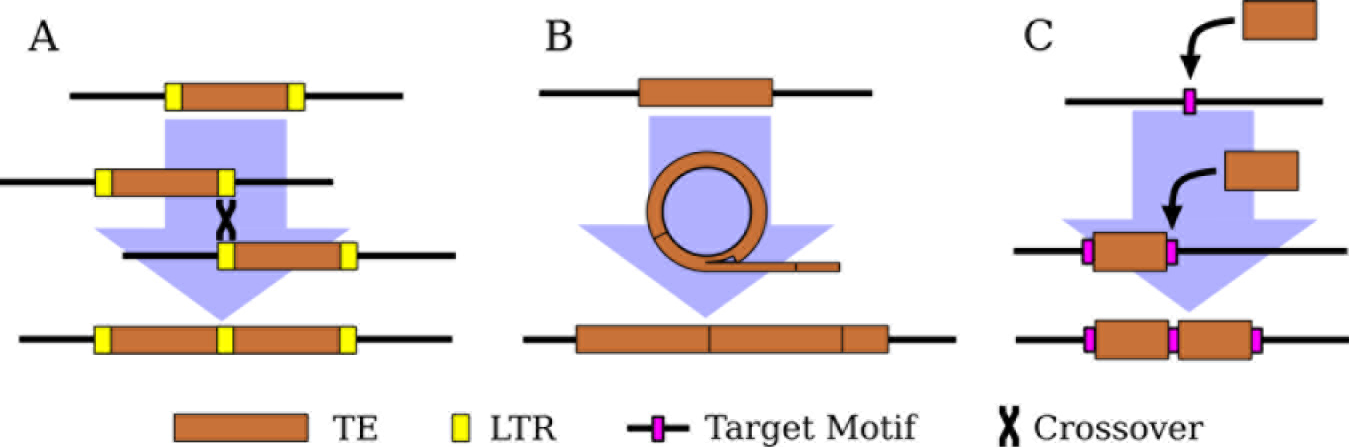
Three mechanisms of tandem TE formation. A) Ectopic recombination between long-terminal repeats (LTRs; shown in yellow) generates tandem LTR retrotransposons with shared LTRs. B) Circularization and rolling circle replication of a TE, followed by insertion of the resulting concatemer. The possible mechanism(s) of circularization remain unclear. C) Two insertions of a TE at the same target site (shown in magenta). Note the preservation of the target site within the tandem junction.

Yet, TEs without intrinsic repeats also form tandem arrays of complete elements (Caizzi et al., 1993) (Miller et al., 1992). One proposed mechanism is rolling circle replication (RCR) wherein an element is circularized and then replicated to form a concatemer that is subsequently reinserted into the genome. (Marsano et al., 2003; Meštrović et al., 2015) (Fig 1B). Alternatively, double insertion of the same element into a single site is possible for TEs that create target site duplications upon insertion. One example is a tandem array of the non-LTR retrotransposon *R1* on the X chromosome in *D. melanogaster* (Kidd and Glover, 1980; Peacock et al., 1981). *R1* has the unusual property of only inserting at a specific site in the multicopy ribosomal RNA genes (rDNA), and the tandem elements are separated by identical 33-nt duplications of rDNA sequence. This pattern is consistent with the tandem originating when two *R1* elements inserted in the same rDNA unit and then subsequently expanded by unequal exchange (Roiha et al., 1981) (Fig 1C). Tandem dimers of DNA transposons have been found in bacterial genomes, and these also contain target-site duplications between the tandem elements (Dalrymple, 1987; Prudhomme et al., 2002), hinting that double insertions may not be limited to elements with insertion site preferences as extreme as in R1.

Whatever the generative process may be, that there are two examples of TEs transitioning to satellite sequence in *D. melanogaster* (Caizzi et al., 1993) (Kidd and Glover, 1980) suggests that it occurs frequently enough that its early stages may be detectable in a survey of population variation. A few tandem TEs, mostly LTR retrotransposons or complex nested insertions, were identified in analyses of the genome assembly (Bergman et al., 2006; Kaminker et al., 2002). However, a full assessment of the mechanisms and frequency with which TEs generate tandem arrays remains unexplored.

Largely this is due to the wider challenge of applying the most comprehensive population genomic resource available—short-read Illumina data—to investigating the evolutionary dynamics of repetitive DNA. One recent analysis of simple satellites (2-10 bp long) across populations of *D. melanogaster* identified considerable variation and population differentiation for a number of satellites (Wei et al., 2014), but similar surveys for complex satellites are more challenging. However, the existence of TE-derived satellites provides a potential opportunity: rather than searching for the emergence of satellites from all possible single-copy sequences, tandems arising from repeats that are normally dispersed rather than tandemly arranged, might yield a tractable model for studying the early stages of satellite evolution.

To identify tandem structures formed by known repeats and ask how these vary across individuals, we developed a bioinformatic pipeline, ConTExt, that utilizes paired-end next-generation sequencing (NGS) data. Applying our pipeline to the Global Diversity Lines, a published panel of 85 *D. melanogaster* strains from five populations (Grenier et al., 2015), we find that TEs frequently form tandem dimers by inserting multiple times at the same locus, continuously generating new substrates of sequences from which satellite arrays can arise.

## RESULTS

### ConTExt identifies repeats from paired-end Illumina data

Paired-end reads are powerful for detecting the junctions arising from structural rearrangements in unique sequences such as deletions, inversions, translocations, and tandem duplications (Bashir et al., 2008; Rogers et al., 2014). Conceptually the problem of identifying the junctions of structures formed by repeats is identical, but is complicated by the fact that repeat-derived reads can rarely be mapped to a unique locus in the reference genome. However, such reads can generally be uniquely mapped to a specific repeat family, and this property has been successfully leveraged in several bioinformatic tools to identify TE insertions into unique sequence (Hormozdiari et al., 2010; Kofler et al., 2012). We extend this idea to identify all types of junctions involving repetitive sequence, including insertions into unique and repetitive sequence, deletions and inversions internal to a repeat, and tandem duplications.

Aligning to the set of all individual repeats present in a reference genome provides the power to detect reads originating from highly divergent variants, but results in reads being distributed across many different sequences. On the other hand, aligning reads to repeat consensus sequences is less powerful in detecting divergent copies, but organizes all reads from a repeat family in the same place, greatly simplifying visualization and downstream analyses. We therefore combine these two approaches into a single pipeline (Fig 2A). In trial runs, we recovered about 20% more repeat derived reads using this two-step procedure than when we aligned only to consensus sequences, with most of the increase coming from relatively ancient repeats such as *INE-1*. We also aligned reads to the repeat-masked reference genome, allowing the detection of junctions between repeats and unique sequence.

**Figure 2:**
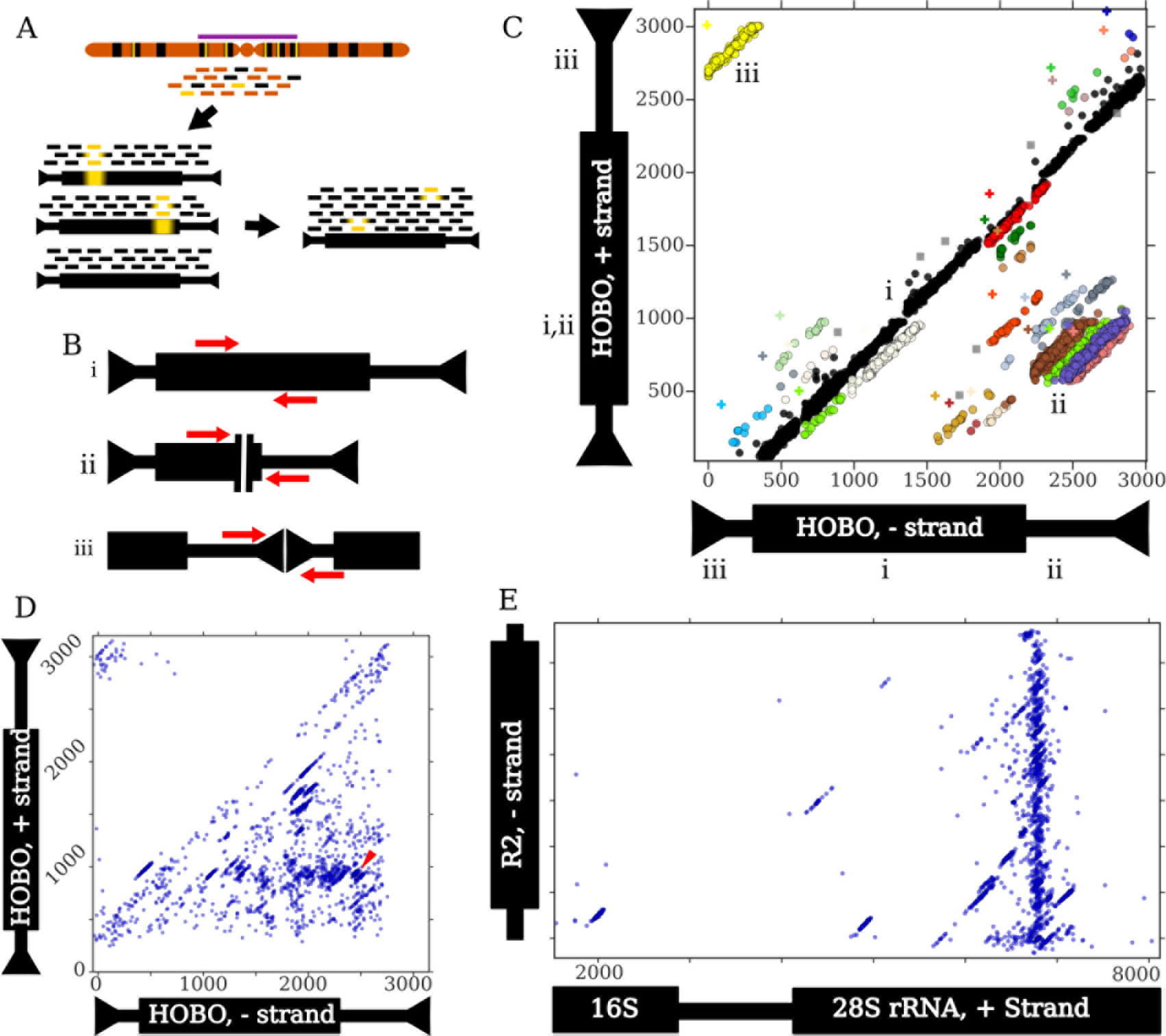
An outline of the ConTExt pipeline and examples of identified structures. A) Sequencing reads are derived from genomic DNA, which has many copies of a particular repeat family (black) dispersed among single-copy sequence (orange); some repeat copies contain divergent regions relative to the consensus (yellow bars), particularly those in the pericentromeric heterochromatin (purple bar). The repeat-derived reads are aligned to a set of individual repeats identified in the reference genome, which includes divergent elements to provide increased power to align reads. Alignments to these individual elements are then collapsed onto a consensus sequence for that repeat family. Inverted arrowheads indicate short terminal inverted repeats (TIRs) that are common to many DNA transposons. LTR retrotransposons have longer direct repeats at their termini. Thin and thick bars represent non-coding and coding sequences, respectively. B) Schematics of paired-end reads spanning i) a region of sequence concordant with the consensus, ii) the junction of an internal deletion, and iii) the junction of a head-to-tail tandem of a typical DNA transposon. C) Paired-end read alignments from strain I03 to the Hobo element represented as a two-dimensional scatterplot. Each dot represents the 3’ end of a read, and its position on the X- and Y-axes corresponds to its aligned position on the minus and plus strands of the Hobo consensus, respectively. Black dots denote concordant reads, other colors correspond to clusters identified by the EM algorithm, and grey squares are reads flagged as possible artifacts. The plus symbols indicate the estimated junction location for the cluster with the corresponding color. The thick black box on the Hobo schematic represents the coding sequence (CDS). The Roman numerals indicate how the three types of structures shown in B correspond to patterns in the scatter plot and where the reads map on each of the axes. i) Concordant reads which form the central diagonal, ii) reads spanning internal deletions, iii) reads spanning head-to-tail tandem junctions. Note that read pairs where both ends map to the same strand (e.g. head-to-head tandems) require a different scatterplot to detect. D) A scatter plot representing the distribution of junctions involving Hobo across all GDL strains. Each dot represents a junction estimated from an identified cluster. The red arrowhead indicates the location of the deletion identified previously in the Th Hobo subfamily (Periquet et al., 1994). E) A scatter plot depicting all junctions between the minus-strand of the R2 retrotransposon and the plus-strand of rDNA. The thick black bar on the rDNA schematic represents the transcribed rRNAs, the thick black bar on the R2 schematic indicates its CDS. The first ~1,500 bp of the rDNA cistron is not shown because only a few low-frequency R2 junctions are present there. The plot successfully identifies that most R2 insertions occur at the same position in the 28S rDNA subunit, as previously demonstrated (Kojima and Fujiwara, 2005; Stage and Eickbush, 2009).

Once reads are organized relative to consensus sequences, we consider the alignment patterns to identify junctions (Fig 2B) and use mixture modeling to cluster and resolve the many junctions that map to each consensus (Fig 2C, Fig S1D). This clustering strategy had >97% recall for junctions supported by at least three reads (Fig S3C) and the ability to resolve nearby junctions was consistent across the GDL, speaking to the uniformity of the sequencing library preparations (Fig S3A,B, Supp Table 1). Once we identify these clusters, we estimate the underlying junctions (Fig 2C) and visualize their distribution across all samples in the dataset (Fig 2D). We cannot accurately infer tandem structures that contain intervening sequence larger than the insert size of the sequencing libraries (on average 338 bp), as we will only detect the junctions between the elements and the intervening sequence, not the elements themselves; this limitation applies mainly to tandem LTR retrotransposons that have large LTRs.

Our pipeline detects various structures involving repeats, including tandem junctions (Fig 2B, C), insertions into unique and repetitive sequence (Fig 2E), and internal deletions (Fig 2B, C). While we focus here on tandems, we note that we successfully identified known internally deleted elements, such as the *Th hobo* variants (Fig 2D) (Periquet et al., 1994) and the *KP* nonautonomous *P* element (Black et al., 1987). We also identified known nested repeats such as *R*-element insertions into the ribosomal RNA genes, including both full-length and distinct 5’-truncated insertions (Fig 2E).

### Transposable elements of all three major types frequently form tandems

Most transposable elements can be detected in tandem in at least one strain but the three major types of TEs show distinct patterns of tandem junctions. (Figure 3; Fig S5). We divide tandems into three types, with head-to-tail tandems being likely to involve full-length elements and/or have intact termini, while tail-to-internal and internal-to-internal tandems are likely to involve 5’-truncated or internally deleted elements (see Fig. 3 for additional interpretations). We do not consider tandems in an inverted orientation, as they cannot expand by unequal exchange and so are unlikely to give rise to satellite sequence.

**Figure 3:**
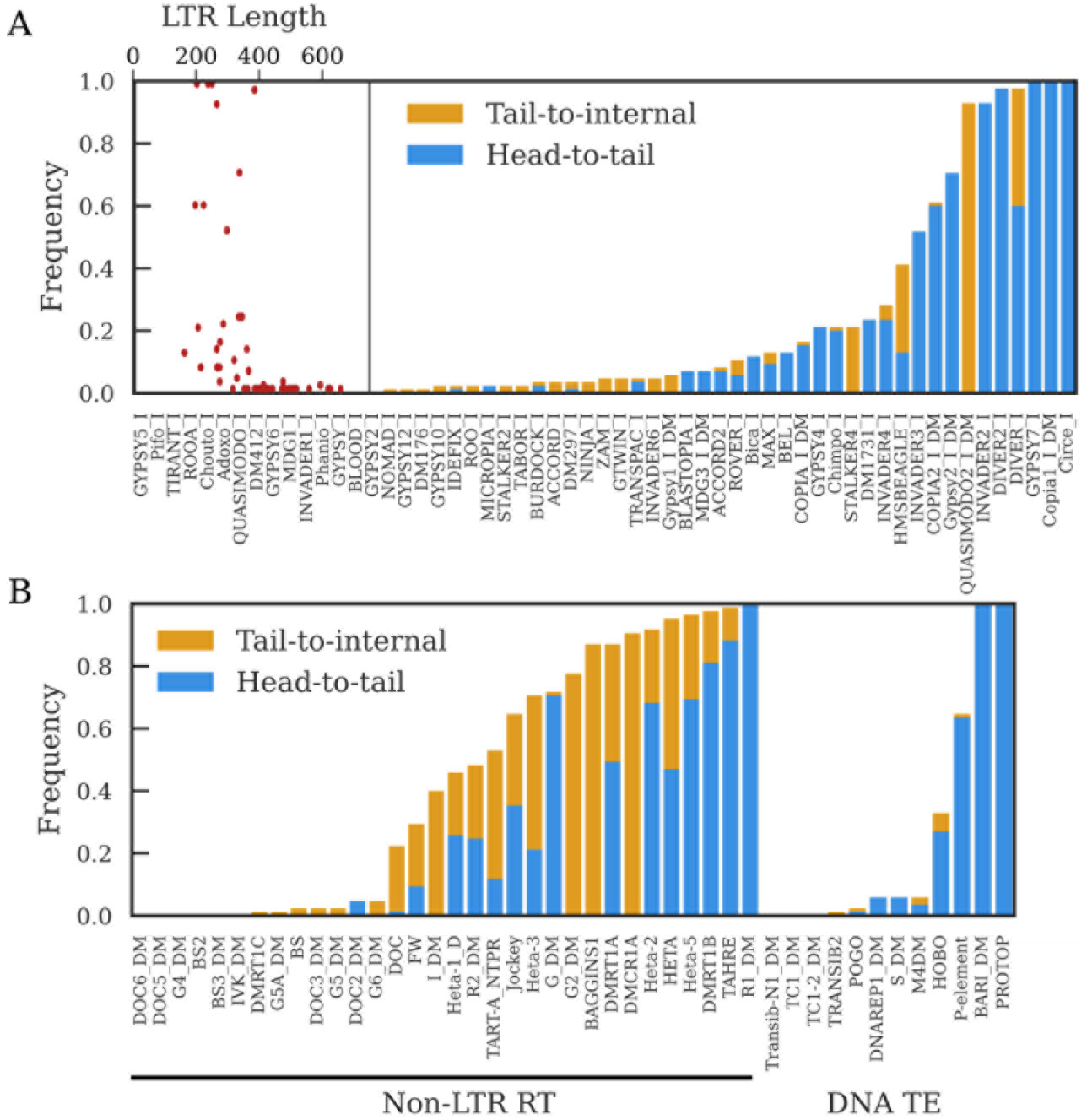
The proportion of GDL strains in which a tandem junction was identified for A) LTR retrotransposon families and B) non-LTR retrotransposon families and DNA transposon families. Head-to-tail tandems have junctions involving the first and last 200 nt of the consensus sequence. Tail-to-internal junctions have junctions between the last 200 nt of the consensus sequence and internal sequence; these are consistent with tandems involving 5’-truncated elements, though they can also be formed by nested insertions. We do not depict the frequency of internal to internal tandems because they are present in most strains, but generally at low copy number; SF5 provides a more informative visualization of internal-to-internal tandem variation. The scatter plot inset in A) depicts the relationship between LTR length and the frequency of detecting head-to-tail tandems for each LTR retrotransposon family.

#### LTR retrotransposons

LTR retrotransposons have a high propensity to form tandems because they are flanked by direct repeats which provide a substrate for tandem formation, with recombination between LTRs yielding structures where adjacent TEs share an LTR (Fig. 1A) (Ke and Voytas, 1997). We detect the majority of LTR retrotransposons in tandem (Fig 3A), though many involve internal sequence and are present at low-copy number (Fig S5A). These internal-to-internal tandems are consistent with deletions that span the junctions of head-to-tail tandems. We less frequently observe head-to-tail tandems, likely because we have limited power to detect tandems when the LTR is longer than the average gap size (~330 nt) (Fig 3A). Notably however, all LTR retrotransposons with LTRs shorter than 300 nt are detected as head-to-tail tandems in at least one strain. Given this and the abundance of internal-to-internal tandems, we conclude that most LTR retrotransposons frequently form tandems.

#### Non-LTR retrotransposons

Unlike LTR retrotransposons, non-LTR retrotransposons and DNA transposons do not provide their own substrates for unequal exchange, yet most can be detected as tandems, demonstrating that additional mechanisms allow tandem formation (Fig 3B). These include the *Drosophila* telomeric TEs which form head-to-tail tandems whenever two elements of the same family insert consecutively at the same telomere (George et al., 2006). Non-LTR retrotransposons are prone to 5’-truncation due to incomplete reverse transcription during transposition, and consistent with these tandems arising through consecutive insertion events at the same telomere, many telomeric TE tandem junctions are tail-to-internal (Fig 3B). Most other non-LTR retrotransposons also can be detected as tail-to-internal tandems in at least one strain, suggesting that transposition is a widespread process generating tandems among non-LTR retrotransposons (Fig 4A, B, E).

**Figure 4:**
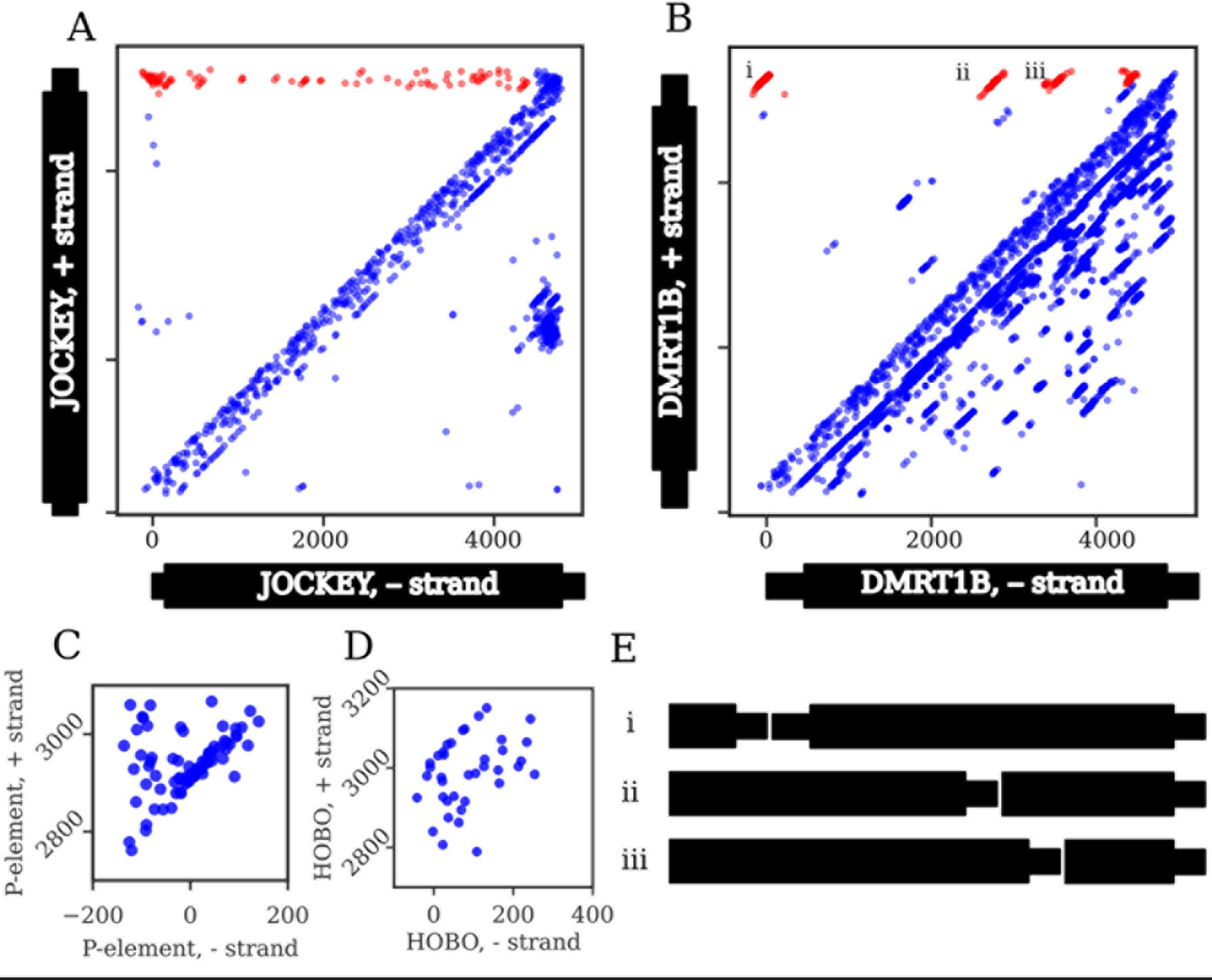
Junction distributions from all strains in the GDL for two non-LTR retrotransposons (A,B) and two DNA transposons (C, D). Note that C and D only show head-to-tail tandem distributions and thus the axes only include the terminal regions. Each dot represents a junction identified from a single strain. If a junction is present in more than one strain, it will generate a diagonal distribution around the true junction coordinate due to estimation errors that arise from the diagonal read pair distributions from which the junction locations were estimated. Head-to-tail and tail-to-internal tandem junctions are highlighted in red for A and B; all junctions are C and D are head-to-tail. The distribution of tandem junctions of jockey (A) are dispersed, with few distinct diagonal clusters, indicating that most individual tandem junctions are low-frequency. By contrast, the four distinct diagonal clusters of DMRT1B (B) indicate junctions at moderate to high population frequency, suggesting that they represent older tandems. C) For the P-element, most junctions fall within a single tight diagonal cluster, consistent with their representing tandem P-elements separated by an 8-bp target site duplication. Several junctions are dispersed above this cluster, consistent with additional sequence of variable length within the junction. D) In contrast, only a few Hobo junctions form a tight diagonal cluster, while most are dispersed below the cluster, consistent with small internal deletions spanning most of the tandem junctions. E) Schematics of the head-to-tail and tail-to-internal DMRT1B tandems denoted with Roman numerals i-iii in B.

The population frequency of tandem junctions across all strains provides insight into the dynamics of tandem TE formation (Fig. 3). For *TAHRE* and *jockey* we find junctions between the 3’ end (i.e. tail) and many different internal locations (Fig 3B, 4A), and these junctions are at low frequency (between 1/85 to 7/85 strains) (Fig 4A). These results strongly indicate many recent and independent tandem forming events. In contrast, *DMRT1B* shows tail-to-internal junctions involving four distinct internal truncations (Fig 4B), found at intermediate frequencies (between 7/85 to 40/85 strains), indicating four independent events that occurred further in the past than the *jockey* and telomeric TE tandems, and that DMRT1 tandem formation subsequently ceased. Taken together, these results suggest that non-LTR tandems are regularly generated within *D. melanogaster*, but by different elements at different periods of time.

#### DNA transposons

DNA transposons are primarily detected as head-to-tail tandems (Fig 3B, Fig S5C), but the pattern of junction estimates suggests that small deletions frequently span the tandem junctions. The junction estimates for *P*-element dimers form a tight diagonal distribution, suggesting recently formed dimers with intact termini (Fig 4C). For *hobo*, an older resident of the *D. melanogaster* genome, we observe a more diffuse distribution, consistent with small deletions near or spanning the tandem junction that are specific to each genome (Fig 4D).

### Multiple insertion drives rapid formation of TE tandems during periods of active transposition

P-element tandems are particularly informative of the dynamics and mechanism of tandem formation because the P element swept through *D. melanogaster* populations during the mid-20^th^ century (Bingham et al., 1982; Engels, 1992; Kelleher, 2016). Given this recent invasion, it is striking that we find tandem P-elements in over half of the GDL strains, indicating that tandem TEs form rapidly during periods of high transpositional activity (Fig 5A). Head-to-head P-element tandems were frequently generated during a genetic screen (Tower et al., 1993), but the majority of strains we analyzed harbor head-to-tail (55/85) rather than head-to-head tandems (8/85). This discrepancy may be due to selection removing head-to-head tandems from natural populations over short timescales, as long inverted repeats are prone to forming cruciform DNA secondary structures (Leach, 1994). Alternatively, this may reflect technical bias, if amplicons containing head-to-head tandem junctions form hairpin secondary structures that decrease their PCR amplification efficiency.

**Figure 5:**
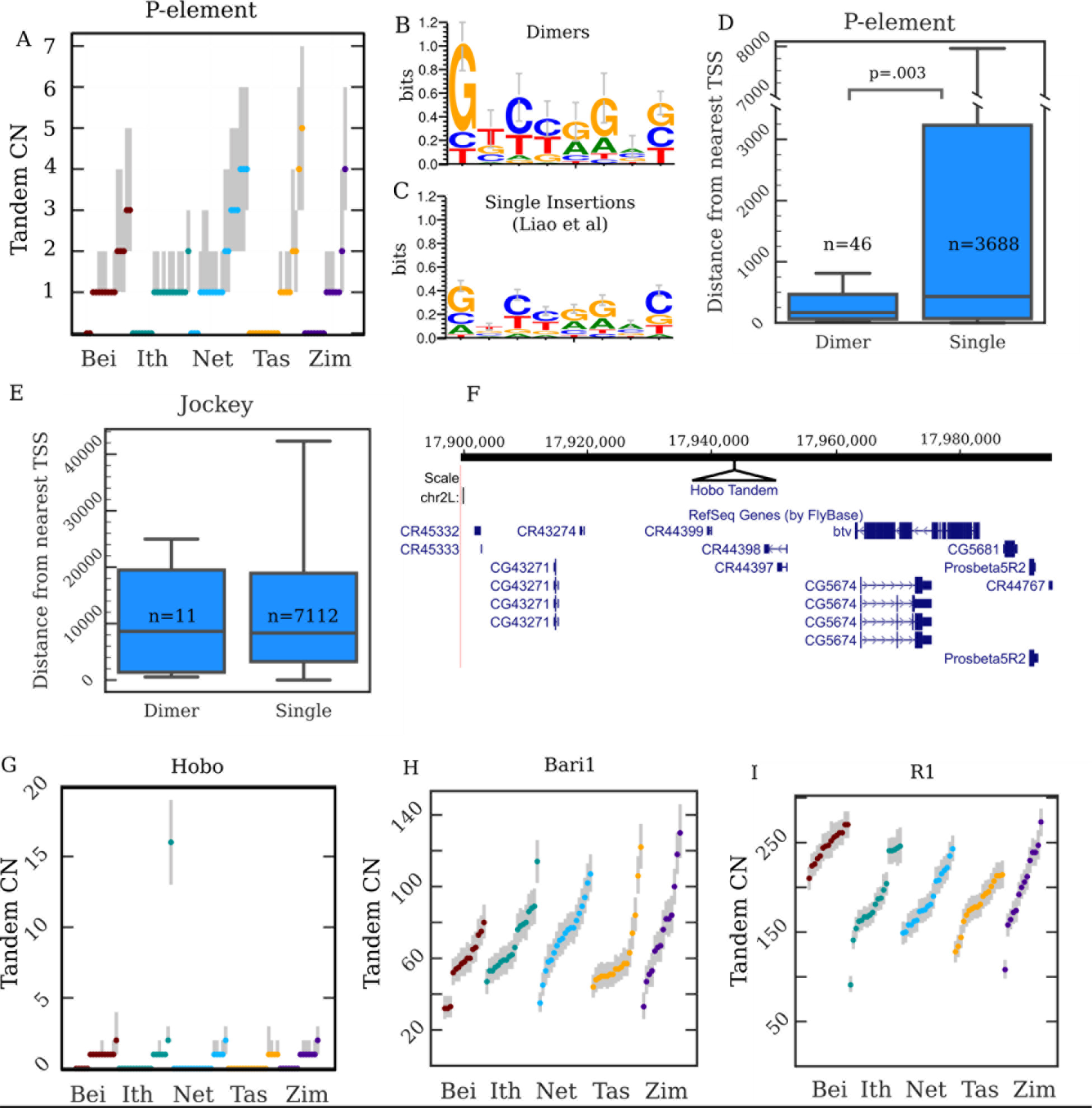
Copy number, location, and sequence of TE tandem junctions. A) Copy number (CN) distributions for the P-element. The dots are maximum a posteriori estimates in a particular strain while the grey lines indicate corresponding 98%-credible intervals. B, C) Sequence Logos constructed from the 8-nt motifs we found within the junctions of P-element tandem dimers (B) and that Liao et al. (2000) reported for P-element TSDs (C). D) A boxplot depicting the distances to the nearest TSS from P-element dimers and single insertions. The numbers within the plots indicate the number of insertions in each category. The p-value is for a Kolmogorov-Smirnov test. E) A similar plot but for Jockey elements. There is no significant difference between Jockey singles and dimers. F) A UCSC genome browser view of the region on chromosome 2L inferred to contain the hobo tandem array in strain I03, with the site of the hobo tandem added in as a black triangle. We mapped the hobo insertions to position 17,943,032. G-I) Copy number (CN) distributions for hobo (G), Bari1 (H) and R1 (I) tandems. The dots are maximum a posteriori estimates in a particular strain while the grey lines indicate corresponding 98%-credible intervals.

To identify the mechanism generating P element tandems, we reasoned that if TE tandems are driven by multiple insertions at the same site, the tandem junction should contain a target site duplication (TSD) of the insertion site (Fig 1C), as has been observed in bacterial DNA transposon dimers (Dalrymple, 1987). We examined reads containing fully intact head-to-tail junctions and found that the majority contain 8 nt of intervening sequence, the same length of the known *P* element TSD. We generated a consensus motif (GTCTAGAG) and found that it is nearly identical to the TSD consensus motif previously identified from 1,469 single *P* element insertions (Fig 5B, 5C) (Liao, 2000). We conclude that *P* element tandems are formed by double insertion at the same site. Importantly, the motifs found at tandem junctions have a higher information content at each position than single-insertion sites, particularly at the first two nucleotides, (sign test, *p*=*.008*), suggesting that *P* element tandems are more likely to form at sites that more closely match its preferred target sequence.

### Genomic distribution of tandem dimers

The target site duplications found in many tandem dimers originate from the locus into which the TEs inserted, and thus contain information about the location of the dimer. We reasoned that we should be able to infer the location of some dimers by identifying every instance of the TSD in the reference genome and asking which ones also contain evidence of TE insertion at that site in that GDL strain (Fig S6 A-F). Because TSDs are short and contain limited information, we imposed a number of filtering steps to restrict ourselves to dimers that could be confidently mapped to a single locus (see Methods).

We successfully mapped 72 dimers, 47 of which are euchromatic using the heterochromatin boundaries defined by (Riddle et al., 2011) (Supp Table 2). *P*-elements comprise the majority of mapped dimers (46/72 mapped dimers), followed by *jockey* elements (11/72). *P*-elements dimers are significantly closer to the transcription start sites of genes (median distance = 169 bp) than are single insertions (median distance = 430 bp) (Fig 5D). Because *P*-elements preferentially insert near the promoters of genes (Spradling et al., 2011), the enrichment of dimers near transcription start sites supports the idea that dimers form at strong insertion sites. This is consistent with the higher information content we observed at TSDs within *P*-elements dimer junctions (Fig 5B). Indeed, several of the tandems we mapped are adjacent to genes previously identified as among the strongest *P*-element insertion hotspots: *apt, RapGAP1*, *Hers*, *Hsromega*, *Men*, and *mir-282* (Spradling et al., 2011). Furthermore, we identified dimers near the transcription start site of *Hers* in three strains (I26, N17, and T29A) and adjacent to *Hsromega* in two strains (B11 and B23), all containing different target site duplications, strongly suggesting that they formed independently. The distance between the 11 *jockey* dimers we mapped and the nearest gene was comparable to that of single insertions, indicating again that the contrasting result with *P*-elements reflects its insertion site preference near promoters (Fig 5 D, E). Moreover, among the ten mapped *jockey* dimers where the 5’-ends of both elements could be identified, six dimers involved clearly distinct 5’-trunctations (>500 nt difference), further supporting our conclusion that these dimers arise by double insertions (Fig S6A-F).

### TE dimers can expand into larger arrays

The abundance of TE tandems that we discovered potentially provides the substrate for expansion by unequal exchange. We therefore searched for larger TE tandems that may be polymorphic among the GDL populations, and discovered one such expansion for the DNA transposon *hobo*. Most strains contain no or only one *hobo* tandems but Ithacan line I03 has an estimated 13-19 tandem copies (Fig 5G, 2C). To determine if this represents multiple independently formed tandems or a single expanded tandem array, we again searched all lines for reads containing fully intact head-to-tail tandem junctions. Only four strains contained fully intact head-to-tail *hobo* tandems, consistent with the distribution of junction estimates which suggested that many *hobo* tandems involve elements with deleted terminal sequence (Fig 4F). Uniquely in I03, we found many reads containing an identical 8-nt motif (GTGGGGAC) between the TIRs of the tandem *hobos*. Using the mapping strategy outlined above, we determined that there is only one locus in the I03 genome that contains both the 8 bp motif and a *hobo* insertion, demonstrating that the *hobo* tandem array is found on 2L at position 17,943,032 of the reference, well outside the pericentric heterochromatin and approximately 19.5 kb away from the protein coding gene *Beethoven* (Fig 5F). I03 is the only strain containing a *hobo* insertion at this position, indicating that the tandem likely formed from a recent *hobo* double insertion. Together, these observations strongly suggest that all elements of the array descend from a single tandem dimer. Multiple independent insertions would instead likely involve distinct motifs unless *hobo* has an extremely specific insertion motif. But, if that were the case, we would observe multiple sites in *I03* with this motif harboring *hobo* insertions, which we do not.

### Copy number variation in TE-derived satellites

Expansion events like we observed with *hobo* can eventually give rise to very large arrays and become fixed. ConTExt successfully identified the two known TE-derived satellites in *D. melanogaster*, which we further investigated to understand the dynamics of established satellites. One satellite is comprised of tandemly arrayed copies of the 1.7 kb DNA transposon *Bari1* and is located in two blocks, with the majority of copies in the pericentromeric heterochromatin of the right arm of chromosome 2 (Caizzi et al., 1993; Marsano et al., 2003; Palazzo et al., 2016). The second block is nested in a *Stalker4* LTR retrotransposon residing on an unmapped scaffold (JSAE01000184); the presence of rDNA units and R elements on the scaffold led Palazzo et al. to suggest it may reside in the X or Y heterochromatin (2016). We identify both expected junctions between *Bari1* and *Stalker4* in the all-female GDL sequences (Supp Table 3), indicating that the smaller array cannot reside on the Y chromosome. Previous analyses identified the *Bari1* tandems in all strains examined (n=10) and estimated its copy number at ~80 repeat units (Caggese et al., 1995). We find the *Bari1* array in all 85 GDL strains, ranging from 32 to 130 copies, which equals ~54,000 to 220,000 base pairs in length (Fig 5H).

The second known TE-derived satellite is comprised of R1 elements, which generally insert only at a specific site in the 28S ribosomal RNA gene. Evidence for this satellite first came from a large-insert clone containing five tandemly arrayed R1 copies. They were mapped to the X heterochromatin but inferred to not be directly within the rDNA array, and all had polymorphisms relative to each other (Kidd and Glover, 1980; Peacock et al., 1981). Subsequent analyses suggested that the copy number of the array might be much higher than five but the overall size and organization of the array was not known (Eickbush and Eickbush, 1995; Stage and Eickbush, 2009). We found that the array is enormous and fixed in the GDL, ranging from 91 to 273 head-to-tail tandem junctions, which suggests it ranges from ~500,000 to ~1,300,000 base pairs in length (Fig 5I). Intriguingly, the scaffold containing the smaller *Bari1* tandem described above (JSAE01000184.1) ends with five tandem R1 elements, and the junction between the first R1 and an rDNA unit is clearly evident. We suggest therefore that this scaffold also contains the boundary of the megabase-sized R1 array.

In addition, we find that the type of tandem dimers that gave rise to the R1 array are still being generated. First, we discovered several low-copy R1 tandems that have rDNA TSDs longer or shorter than 33-nts. These are likely dimers that are independent of the R1 array because we confirmed the previous observation that the array is comprised of dimers with a 33-nt TSD of rDNA sequence. Second, we find many strains also contain 5’-truncated tandems (Fig S7B), suggesting the continuous production of R1 dimers distinct from those in the large array. We conclude that our population survey captures R1 elements in both nascent and fixed arrays.

### The R1 array is more heterogenous than the Bari1 array

In addition to copy number variation, we discovered many R1 junctions corresponding to internal deletions, internal-to-internal tandems, and TEs inserted into R1 elements, many of which might reside in the tandem array. Theory predicts that evolution by unequal exchange can organize such structural variation into higher order repeats (HORs) (Stephan, 1989). HORs are features of known satellites, including the centromeric human alpha satellites, and variant higher order repeats can have elevated rates of chromosome missegregation (Aldrup-MacDonald et al., 2016). Thus, given the amount of potential structural variation within the array, we wished to determine whether the array shows evidence for HORs. We could not determine the exact organization of the array due to the impossibility of assembling from Illumina data, but we reasoned that if a junction is interspersed throughout the array its copy number should correlate with the overall size of the R1 array across lines. Because this requires comparing the copy number distributions of specific junctions rather than general categories of structures as we have done above, we needed a principled strategy for matching estimated junctions across strains. To this end, we employed a fuzzy C-means-like algorithm to match junctions across strains, using the uncertainty around each junction estimate to inform cluster assignments (see Supplemental Methods, Fig S4E). We then assessed each junction for a copy number correlation with the head-to-tail tandem junction, determining significance with the Benjamini-Hochberg procedure at a FDR of 1%.

Using this approach, we found 92 junctions involving R1 to have significant positive correlations with array size (Supp Table 3). Neither negative correlations or any of the many junctions corresponding to R1 insertions in the rDNA were identified as significant, either of which would likely reflect false positives, suggesting that technical bias is rare. Among the 92 junctions we found 20 tandems junctions and 24 internal deletions (Fig S7B). One of these tandem junctions is a high copy junction consistent with a small deletion near or spanning the head-to-tail tandem junction, which is present in about one quarter of tandem R1 units; an examination of ISO-1 PacBio reads confirms its presence in the array (Fig S7A) (Kim et al., 2014). Among the 92 we also found a number of TE insertions into R1 elements that are correlated with the copy number of the array. The highest copy examples are an *FW* insertion which expanded to an average of 18 copies, a *Circe* insertion present at an average of 6 copies, and an *Accord2* insertion present at an average of 6 copies (Supp Table 3). The presence of *Circe* within the array is consistent with previous cytological observations (Losada et al., 1999). The copy numbers and degrees of positive correlation we observe also indicate that these structures are dispersed throughout the entire array or constitute subarrays that may be arranged as higher-order repeats, as expansion and contraction events that change the array’s copy number also alter the copy number of these junctions. For comparison, we looked for junctions involving *Bari1* that were correlated in copy number with the *Bari1* head-to-tail tandem junction, and found only 8 junctions, none of which had amplified to multiple copies in any strain (Supp Table 3). The reference genome indicates a Max LTR retrotransposon is inserted into the *Bari1* array, but we find no evidence of the insertion in the GDL strains, suggesting it is specific to the reference strain (Hoskins et al., 2015). Taken together, these observations suggest that the R1 array but not the Bari1 array is heterogenous with respect to deletions and TE insertions, some of which may be arranged into higher order repeats.

## Discussion

### ConTExt successfully identifies repetitive structures in NGS data

Leveraging NGS population genomic datasets to learn about highly repeated sequence is a challenging problem. We employed an alignment strategy that maps repeat-derived reads to repeat consensus sequences and used mixture modeling to interpret the alignment patterns. Applying this method to a panel of five populations, we observed multiple stages of TE-derived satellite evolution on-going within a single species. We successfully detected previously known tandem structures, including tandem junctions amongst all telomeric TE families and large tandem arrays of the *Bari1* and *R1* elements. We also identified internally deleted TEs as well as nested insertions (Figs 2D, E). Using only short-read data, we gained considerable insight into the organization of tandem TEs, highlighting the large amount of information about these understudied structures already available in the thousands of publicly available datasets. ConTExt can thus address a wide variety of other questions about repeat evolution in any species with a genome assembly and repeat annotations.

Our strategies have some limitations imposed by our reliance upon sequence alignments. Repeat-derived reads rarely align uniquely to the reference genome, which means that we cannot locate most of the structures we identify. Further, reliance on consensus sequences to identify repeat-derived reads limits our survey to known repeat families. However, as the TE families in *D. melanogaster* are well-characterized this is unlikely to have strongly biased our analysis. For less well-characterized species, tools exist which can extract repeat consensus sequences out of NGS reads (Novak et al., 2013).

Structure inference from paired-end alignments also has limitations. First, we cannot detect junctions containing intervening sequences longer than the mate pair distance of the library, such as LTRs exceeding 338 bp. Second, chimeric inserts in paired-end data can produce spurious structure discovery calls. To account for this, we only considered structures supported by multiple read pairs with distinct coordinates. Furthermore, it is unlikely that the general patterns we observe are false positives, as false positives should be dispersed across the consensus sequences rather than concentrated in biologically plausible patterns specific to TE types. Indeed, many of the junctions involving non-LTR retrotransposons involve 5’-truncated elements as expected based on previous studies. Moreover, the tandems we find for LTR elements are largely restricted to LTRs below the length cutoff expected based on the library insert size. If false positive were being generated by either mapping artifacts or by random chimeric structures formed during library preparation, we would instead expect to find head-to-tail dimers of LTR retrotransposons regardless of LTR length.

### Transposition drives continuous production of tandem formation

We discovered that the processes by which TEs transition to satellites are actively ongoing in *D. melanogaster*. Tandem TE dimers from which large satellite arrays can expand are common, with multiple TE dimers present in most *D. melanogaster* genomes. The frequency with which we detect LTR retrotransposon tandems is inversely associated with LTR length, which is consistent with the tandem junctions containing LTRs. Such tandem junctions are the likely consequence of ectopic recombination between LTRs, as has been previously observed (Ke and Voytas, 1997). In contrast, several observations strongly suggest non-LTR retrotransposon and DNA transposon tandems are formed by multiple insertions at the same locus. First, it is well documented that repair mechanisms generate direct repeats flanking most TE insertions (Craig, 1997). Thus, a tandem formed by double insertion should contain a duplicate of the target site within its tandem junction. This prediction is borne out by many of the tandems that we identified, in particular the majority of *P*-element dimers harbor intervening sequence matching the element’s known target motif. Second, the patterns of 5’-truncated tandems we observe in non-LTR retrotransposons support multiple insertion events. Non-LTR retrotransposons are prone to losing sequence from their 5’ ends during integration, and most non-LTR retrotransposon tandem junctions we found are between the intact 3’ end of one element and the 5’-truncated end of the adjacent element. We discovered several non-LTR retrotransposon dimers involving elements with distinct 5’-truncations, clear evidence of two independent insertion events.

Third, if TE dimers form through transposition events then periods of high TE activity should also have high rates of dimer formation. Indeed, despite only invading the species in the last century, we find that most strains in the GDL contain tandem P-elements. We further note that the population frequency of particular dimers varies among elements, suggesting discrete periods of dimer formation. Thus, bursts of TE activity likely correspond to bursts of tandem formation. We suggest that the presence and population frequency of tandem dimers may be a useful proxy for identifying recently active TE families. Overall, for those TEs with an insertion site preference, the amplification events which form tandem dimers are common and almost guaranteed by the presence of active copies.

We emphasize that the mechanism of transposition ensures dimer formation. Dimer formation requires only that an element inserts preferentially at certain motifs and generates target site duplications. If so then a new TE insertion does not consume its target site but rather preserves it while generating a new one, enabling subsequent insertions at that locus (Fig 1C). The propensity of most TE families to form dimers highlights the degree of insertion site preference: a TE family which inserted at random sequence would almost never be detected in tandem.

### Tandem dimers expand into large arrays

Having discovered that TE dimers are common in natural populations, we suggest that their subsequent amplification is the major mechanism generating TE-derived satellites. We observed one such event, a copy number expansion of a *Hobo* dimer to ~16 copies in a single line (Fig. 5G). We also characterized the previously discovered array of R1 elements. We confirmed earlier suggestions that it may large, finding that it varies between ~530,000-1,300,000 bp in length. Our analysis of the junctions of the R1 array reveals that it likely originated when an R1 dimer formed within an rDNA unit, and then expanded. We further found that many independent deletions and TE insertions occurred within the array subsequent to its expansion, some of which also expanded in copy number. The obvious candidate for such expansions is unequal exchange.

While the R1 and Hobo arrays contain target site duplications clearly indicating that they originated as a dimer, the junctions in the two *Bari1* arrays do not. Instead the tandem *Bari1* elements display several unusual features: they are missing sequence from their terminal inverted repeats, each ends with a partial element at its 5’ edge, and the smaller array is flanked by a ~500-nt TSD inconsistent with *Bari1’s* usual transposition mechanism (Marsano et al., 2003). Marsano et al. (2003) suggested rolling circle replication as an explanation, with TIR sequence being lost during circle formation and the partial terminal elements resulting from utilization of a random cut site due to the incomplete TIRs (Marsano et al., 2003). Alternatively, the smaller array may represent the migration of sequence from the array on 2R to the X chromosome. Indeed, loop deletions within satellite arrays naturally generate circular DNAs which are speculated to facilitate the spread of satellites to new loci (Cohen and Segal, 2009)(Khost et al., 2017). Regardless, we agree with Marsano et al. that rolling-circle replication with mechanistic biases is a plausible explanation of the *Bari1* array’s origin given the absence of TSDs within the junctions.

However, the absence of target site duplication at the tandem junctions does not preclude the alternative possibility that the larger *Bari1* array arose by tandem insertion of two *Bari1* elements, followed by partial deletion of the terminal inverted repeats at the junction, and then expansion by unequal crossing over. Deletions of TSDs are common for DNA transposons, as we observed for most *Hobo* dimers (24/36). The expansion of such a dimer would result in an array where each junction contains an identical deletion, as is found in the *Bari1* array. While distinguishing whether rolling-circle replication or double insertion is the more plausible explanation for generation of the *Bari1* array will require knowledge of the rate parameters of rolling-circle replication, our results indicate that double insertions rapidly generate abundant source material from which tandem arrays can arise.

### Heterogeneity differences between arrays

The R1 array is substantially more heterogeneous than the *Bari1* array, harboring a number of deletions and TE insertions residing within the array. Such organization is typical of many satellite arrays where TEs tend to accrete to the edges of the array (Khost et al., 2017; McAllister and Werren, 1999). One explanation for these differences in heterogeneity is that the R1 array is older than the *Bari1* array. Consistent with the R1 array being relatively old, we find that most of the structures responsible for its heterogeneity are at relatively high population frequency, indicating that they were present when the GDL populations diverged. Alternatively, differences in the recombination rates experienced by the two arrays might account for their structural differences, as simulation studies suggest that array heterogeneity and higher-order structure tend to arise naturally when the rate of unequal exchange (relative to the mutation rate) is low (Stephan and Cho, 1994). A third possibility is that the sequence identity of the *Bari1* array is maintained by purifying selection.

### Tandem persistence and recombination rate

We noted that DNA transposon tandems frequently lacked terminal sequence, which could reflect either retention or mutation bias. In particular, we observed that the *P*-element dimers, which are younger than 100 years old, generally have intact TIRs, while most of the older *Hobo* dimers lack TIRs. Likewise, a large *P*-element related array in *D. guanche* is comprised of elements missing ~100 nt of terminal sequence (Miller et al., 1992), and all elements in the fixed *Bari1* tandem array have incomplete TIRs (Marsano et al., 2003). The terminal inverted repeats at the ends of DNA transposons are endonuclease cut sites, and the presence of a cut site within a tandem should expose it to elevated rates of double strand breaks and unequal exchange. By contrast, tandems lacking intact TIRs will experience a reduced rate of double strand breaks and are likely to experience fewer recombination events over time. Charlesworth et al. proposed that tandem arrays in regions with high recombination rates should be lost more rapidly than those in low recombination regions, shaping the genome-wide distribution of satellite sequence (Charlesworth et al., 1986). More generally, they proposed that this applies to any satellite with features that reduce the rate of unequal exchange. We suggest that differential persistence may also shape the sequence evolution of tandem DNA transposons, with tandems harboring intact TIRs being lost more rapidly. A second possibility is that the palindrome formed by inverted repeats at the tandem junction leads to hairpin secondary structures, which may be prone to deletions.

### Implications of tandem TEs

While the structures we describe are present in most genomes, they cannot be detected by the standard tools for structural variant discovery. They have thus been largely ignored in previous analyses of TE structural variation despite having known biological effects, such as on gene expression. For example, tandem *P*-element transgenes induce position-effect variegation, the strength of which increases with copy number (Dorer and Henikoff, 1994). This is likely because TE insertions are targeted for heterochromatinization by the piRNA pathway (Brennecke et al., 2007) and these silencing marks can spread into nearby genes and regulatory sequences (Lee, 2015; Shpiz et al., 2014). TEs also frequently carry internal regulatory elements that can be recruited into gene regulatory networks and even alter the three-dimensional organization of the genome (Byrd and Corces, 2003; Feschotte, 2008). Loehlin et al recently described synergistic increases in the expression of recently duplicated genes which may result from concentrating regulatory elements (Loehlin and Carroll, 2016). We suggest therefore that future studies on the functional impacts of TE variation should consider whether the insertions in question are single elements or tandemly arrayed. This is particularly important for elements with strong site preferences such as *P*-elements, with insertional hotspots being most likely to harbor tandem structures.

## Methods

### Overview

We approach the problem of making inferences about repeats from NGS data using two main steps. First, we align reads to the consensus sequences of known repeats. Second, we employ a clustering strategy to infer structures from the distributions of discordant read pairs in each library. Specifically, we sought to identify junctions: sequence coordinates that are non-neighboring in the reference genome but which neighbor each other in the sequenced genome (Bashir et al., 2008). We frame our goal as identifying a generative model that explains the observed distribution of aligned read pairs, and we accomplish this through mixture modeling. This general approach not only allows us to identify the presence and copy number of tandem structures, but also deletions internal to repeats, and insertions into both unique and repeated sequence. Importantly, it can be applied to any genome for which the repeat families are known.

### Constructing the repeat index

We downloaded the RepBase repeat annotations (release 19.06) (Bao et al., 2015) for *D. melanogaster* and supplemented these with a set of simple and complex satellites found at <http://www.fruitfly.org/sequence/sequence_db/na_re.dros>. We also included the rDNA sequence from (Stage and Eickbush, 2007), additional TE sequences from (Zanni et al., 2013), and extracted highly divergent variants of the telomeric TE *Het-A* from the reference genome. Further, the *TART* telomeric TEs contain very long terminal repeats, which we removed from the consensus and placed in separate entries, in accord with how LTRs are handled for LTR retrotransposons (Jurka, 2000)

This set of sequences contained redundancies that would have complicated interpreting the alignments; for example, there were over 20 sequences corresponding to the centromeric 359-bp repeat satellite, and two *Protop* variants which simply corresponded to internally deleted elements. To resolve these issues, we manually curated the index by performing all pairwise alignments to identify entries that share considerable homology (see Supplemental Methods for details). *Batumi*, *Foldback* and *Vatovio* are not included in subsequent analyses due to errors in RepBase entries or their propensity for mapping artifacts. While we subsequently refer to the entries in this index as consensus sequences, not all are true consensus sequences; rather, some entries are representative examples. We use the RepBase nomenclature for repeats with the exception of the *Bari* transposon which we refer to as *Bari1*, following Kaminker et al (2002).

### Read preprocessing

We used Trimmomatic for read quality control (Bolger et al., 2014). We removed all sequence within and subsequent to the 5’-most four nucleotide window where the average PHRED quality score was less than 20. We discard any trimmed reads with lengths less than 40-nt. Because we can only detect a junction if it falls in the gap of a read pair, we trimmed any reads longer than 70-bp down to 70-bp from their 3’-ends to increase the size of this gap.

### Aligning reads to repeats

The simplest way to determine from which repeat family a read derives is to align reads to a set of repeat consensus sequences. However, the consensus sequences of TE families frequently resemble the most active subfamily and can be quite diverged from older insertions. Reads originating from insertions of old families, like DINE, often do not align even under very permissive alignment parameters. To circumvent this, we employed a two-step alignment procedure, first using Bowtie2 to align the reads to the set of all individual repeats extracted from the reference genome (including the unmapped contigs), and then collapsing these alignments onto the corresponding consensus sequence.

We first used RepeatMasker (Smit et al., 2015) to both mask release 6.01 of the *D. melanogaster* reference genome (Hoskins et al., 2015) for repeats and to identify the location of all instances of each repeat family, using the most sensitive seed setting. We extracted these repeats from the reference to construct an index of individual insertions. We then used Bowtie2 (version 2.1.0) to align the reads in each read pair as single-end reads to both the repeat-masked reference genome and the index of individual repeats (Langmead and Salzberg, 2012). We used parameters [bowtie2-align -p <# threads> -<phred format of data> --score-min L,0,<-.64 or -1.0> -L 22 -x <Index> -U <infile> -S <outfile>] and use -.64 as the alignment threshold for the reference genome and -1.0 as the alignment threshold for the insertion index.

Alignments were then collapsed onto the corresponding repeat consensus sequences. To accomplish this, we first used BLASTn to align all individual repeats to all repeat consensus sequences and kept those alignments where the identity was at least 80%. If an individual repeat aligned multiple times to the set of consensus sequences, we chose the alignment with the lowest e-value. If there were multiple alignments tied for e-values, we chose the longest alignment. We then used the BLAST traceback strings to convert the aligned coordinates of reads from individual insertion to the corresponding coordinates on consensus sequences. If the new aligned length of the read on the consensus was less than 50% or greater than 150% of the original read length, that alignment to the consensus was rejected and the read was marked as unaligned. We updated the CIGAR and MD strings to reflect the new alignment. We filter out any reads which align to the reference genome with mapping quality scores less than 20. Because the index of TE insertions is highly repetitive, mapping quality scores are not informative of alignment quality and so we do not apply the same filter to repeat aligned reads.

### Estimating the gap size distribution

The distribution of reads spanning a junction depends upon the size distribution of read pairs in the library. We refer to the distance between the 5’ ends of a concordant read pair as the insert size, and the interval of sequence between the 3’ ends of a read pair reads as the gap. For a junction to be detected, it must be spanned by the gap (junctions interrupting a read will likely prevent its alignment), so for each library we estimated the gap size distribution with a kernel density estimate, choosing the bandwidth by 2-fold cross-validation (see Supplemental Methods for more details). For simulations of read distributions, we use the kernel density estimate conditioned on the reads spanning a junction which accounts for small inserts being less likely to span a junction and is approximately given by:

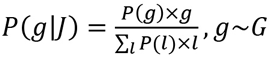, where G is the gap size distribution.

In all subsequent sections, we consider a read pair concordant when its two reads map to opposite strands and are oriented toward each other, and when its gap size falls between the .5%-percentile and the 99.5%-percentile of the gap size distribution.

### Representing paired-end alignments as two-dimensional scatterplots

A read pair can be represented as a point in a two-dimensional space, where the X-axis represents the sequence and strand to which one read maps, and the Y-axis represent the sequence and strand to which the other read maps (Fig S1B,C, Fig 2C). This is an effective visualization strategy for manually examining the patterns of TE insertions into unique sequence and into repetitive sequence. Organizing reads where both ends map to the same sequence requires additional constraints. For read pairs that map to opposite strands of the same sequence, we assign the reverse strand to the X-axis and the forward strand to the Y-axis. For read pairs that map to the same strands of the same sequence, there is ambiguity as to which axes the reads should be assigned. In the case of forward-forward read pairs, we assign the read with the higher sequence coordinate to the X-axis, and for reverse-reverse, we assign the read with the lower sequence coordinate to the Y-axis.

### Discovering structures with mixture modeling

The problem of structural variant discovery can be framed as trying to identify clusters of read pairs that span junctions. While agglomerative clustering strategies are successful at identifying structural variation in unique sequence (Medvedev et al., 2009), the alignment patterns of repeat-derived reads are more challenging to resolve. This is because one is collapsing reads derived from up to megabases of sequence onto consensus sequences less than 10kb in length, and so read pairs representing distinct junctions are often crowded and sometimes interspersed. Mixture modelling, however, provides well-founded tools for clustering data, especially when clusters are partially overlapping. Therefore, we model the distribution of discordant read pairs within a scatterplot with a Gaussian Mixture Model (GMM) (Fig S1D). A junction involving a repeat will generate a distribution of discordant read pairs (Fig S1B, C), and so the set of discordant read pairs, *X*, in a scatterplot can be thought of as arising from a mixture of many distributions, each corresponding to a junction (Fig S1D):

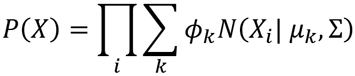

Thus, each component, *k*, in the GMM corresponds to a junction, with the mean, *μ*_*k*_, relating to the junction’s sequence coordinates, the mixing proportion, *ϕ*_*k*_, relating to the number of read pairs spanning that junction, and the covariance, Σ, reflecting the library’s gap size distribution. The actual distribution of read pairs spanning a junction is not Gaussian (Fig. S1C), however the approximation makes the problem tractable and we use Gaussians with sufficiently large covariances to cluster the read pair distributions. We fit the GMM using an accelerated Expectation-Maximization algorithm (Dempster et al., 1977) (Varadhan and Roland, 2008). We then use the fitted GMM to group read pairs into clusters that correspond to junctions, assigning each read pair to the most likely component (Fig 2C, Fig S1D). Once clusters are identified, we remove clusters that are possibly technical artifacts and estimate the sequence coordinates and copy number of the underlying junctions in a manner that accounts for GC-bias in read depth. For further details of the EM implementation, covariance selection, and post-processing of the identified clusters, see Supplemental Methods. Summaries of clustering parameters and performance can be found in Supplemental Table 1.

### Mapping tandem dimers to specific TE insertions

To infer the location of a tandem dimer, we first identified sequencing reads in that strain containing the tandem junction (Supplemental Methods). From these reads we identified any intervening sequence at the junction, and identified every locus in the reference genome matching the intervening sequence. For sequences 9-bp or longer, we used BLASTn with an e-value cutoff of 10 and accepted the top hit and all other hits whose e-values were within 2-orders of magnitude of the highest e-value. For intervening sequences 7- or 8-bp in length, we required an exact match to either the sequence or its reverse complement, and employed string matching. We did not attempt to map any dimer whose intervening sequence was less than 7-bp.

We reasoned that if one and only one matching locus contained an insertion of that TE family, it was the likely location of the dimer. So, we identified the location of every insertion of the corresponding TE family in that strain (Supplemental Methods). We matched each intervening sequence to a TE insertion if the estimated location of at least one of its junctions in the reference is within 150-nt of the intervening sequence. We only considered dimers that could be uniquely mapped to a single location. We note that while we putatively mapped R1 tandems to two locations in autosomal heterochromatin, these were driven by partial alignments with distinct target site duplication, and are likely artifacts considering the high site-specificity with which R1 normally inserts; thus, we excluded R1 from our efforts to map dimers.

### Gene annotation

We downloaded the RefSeq gene annotations for *D. melanogaster* from UCSC’s Table Browser and excluded all computed genes and RNAs (entries named CG#### or CR####).

### Aligning Pacbio reads

We aligned Pacbio reads to the repeat index using BLASTn, using a linear gap penalty of 2 to account for the high rate of indels and imposed an e-value cutoff of .01.

### Categorizing tandem junctions

We divide the tandem junctions we observe into three broad categories based upon their sequence coordinates: head-to-tail, tail-to-internal, and internal-to-internal. We define head-to-tail junctions as those within 200-nt of both the 5’ and 3’ ends. We define tail-to-internal as junctions within 200-nt of the 3’-end, but not the 5’-end. All other tandem junctions are classified as internal-to-internal. We exclude junctions where the coordinates are within 400-nt of each other, as these potentially reflect groups of concordant reads misidentified as discordant. We restrict this analysis to TE families estimated to contribute at least 20kb of sequence to at least one genome, based on coverage of the consensus normalized by GC-corrected read depth.

### Matching junctions across samples

Fitting the GMM to the data allows us to identify junctions within each sample, but for some questions we needed to match these junctions across samples. To do this automatically, we employ a second fuzzy clustering step, that uses the estimated uncertainty around each junction estimate to define cluster membership weights based on the probability that multiple inferred junctions reflect the same structure. For more details, see Supplemental Methods.

